# A circulating T-cell differentiation marker to predict response to immune checkpoint inhibitors

**DOI:** 10.1101/2020.06.13.095844

**Authors:** Takayoshi Yamauchi, Toshifumi Hoki, Takaaki Oba, Vaibhav Jain, Hongbin Chen, Kristopher Attwood, Sebastiano Battaglia, Saby George, Gurkamal Chatta, Igor Puzanov, Carl Morrison, Kunle Odunsi, Brahm H. Segal, Grace K. Dy, Marc S. Ernstoff, Fumito Ito

## Abstract

Immune checkpoint inhibitors (ICI) have revolutionized treatment for various cancers; however, durable response is limited to only a subset of patients. Discovery of blood-based biomarkers that reflect dynamic change of the tumor microenvironment, and predict response to ICI will markedly improve current treatment regimens. Here, we investigated a role of CX3C chemokine receptor 1 (CX3CR1), a marker of T-cell differentiation, in predicting response to ICI therapy. Successful treatment of tumor-bearing mice with ICI increased the frequency and T-cell receptor clonality of the peripheral CX3CR1^+^CD8^+^ T-cell subset that included an enriched repertoire of tumor-specific and tumor-infiltrating CD8^+^ T cells. Furthermore, an increase in the frequency of the CX3CR1^+^ subset in circulating CD8^+^ T cells early after initiation of anti-PD-1 therapy correlated with response and survival in patients with non-small cell lung cancer (NSCLC). Taken together, these data support T-cell CX3CR1 expression as a blood-based dynamic biomarker to predict response to ICI therapy.

---

Cancer immunotherapies that target the immune checkpoints, such as cytotoxic T lymphocyte associated antigen 4 (CTLA-4), programmed cell death protein-1 (PD-1), and PD1 ligand-1 (PD-L1), have transformed the therapeutic landscape of a variety of malignancies^1–3^. However, despite unprecedented and durable clinical responses seen across diverse tumor types, only a fraction of patients achieve durable responses. Moreover, unusual response patterns such as pseudoprogression and delayed response, a dichotomous outcome, potentially severe toxicity, and high cost indicate a critical need for a reliable predictive biomarker^4–7^.

Baseline PD-L1 expression on immune and tumor cells, preexisting infiltrating CD8^+^ T cells, and tumor mutational burden (TMB) correlate with response^2–11^; however, the use of these pretreatment markers are hampered by the significant overlap between responders and non-responders, limited quantity and quality of the tissue, and/or lack of standardization^12–14^. Analysis of serially collected tumor samples could aid in the assessment of evolution of the tumor 15-18 microenvironment (TME) during immune checkpoint inhibitor (ICI) therapy^15–18^; however, this approach is invasive and challenging for visceral tumors such as non-small cell lung cancer (NSCLC). Discovery of dynamic circulating immune biomarkers that reflect the evolution of adaptive immunity in the TME and are early predictors of clinical response to ICI would be of value to guide selection of patients most likely to benefit from ICI therapy.

Emerging blood-based biomarkers such as exosomal PD-L1, TMB and T cell receptor (TCR) sequence in cell-free DNA, and hypermutated circulating tumor DNA associate with response^19–23^; however, these approaches require complex platforms, and/or bioinformatics analysis, that limit their widespread application in community-based clinical practice. Since ICI targets T-cell regulatory pathways, utility of surface and intracellular proteins expressed on T cells have been investigated as a potential biomarker for response^8, 9, 24–28^. Of these, the proliferation marker Ki-67 has been extensively investigated. However, most studies showed Ki-67 expression only transiently increased in subsets of peripheral blood (PB) CD8^+^ T cells after the first cycle with unclear predictive and prognostic value as a stand-alone predictive biomarker for ICI^8, 9 24–27^

Recently, CX3C chemokine receptor 1 (CX3CR1) was found to be a marker of T-cell differentiation, where CX3CR1^+^CD8^+^ T cells were progeny of CX3CR1^-^CD8^+^ T cells, and exhibited robust cytotoxicity in anti-viral immunity^29, 30^. Mechanistically, CX3CR1 is stably expressed on CD8^+^ T cells through unidirectional differentiation from CX3CR1^-^CD8^+^ T cells during the effector phase^30–32^, which theoretically provides an advantage as a biomarker compared with transiently expressed molecules on T cells. Indeed, an increased frequency of PB CX3CR1^+^CD8^+^ T cells has been observed in a few patients who responded to anti-VEGF and anti-PD-L1 antibodies (Ab)^33^ or chemotherapy and anti-PD-1 Ab^34^; however, no studies have evaluated the utility of CX3CR1 on T cells as a blood-based predictive biomarker only for ICI therapy.

Here, we hypothesized that changes in the frequency of PB CX3CR1^+^CD8^+^ T cells would correlate with response to ICI and help identify responders versus non-responders early after initiation of therapy. We investigated the frequency of CX3CR1^+^CD8^+^ T cells in PB before and during ICI therapy, and delineated the TCR repertoire in peripheral CX3CR1^+^CD8^+^ T-cell subsets and CD8^+^ tumor-infiltrating lymphocytes (**TILs**) using preclinical models. To understand the clinical utility of CX3CR1 as a circulating T-cell biomarker, we analyzed longitudinal PB samples from patients with NSCLC undergoing anti-PD-1 therapy and evaluated the predictive and prognostic value of changes in the frequency of PB CX3CR1^+^ CD8^+^ T cells. Our results support circulating CX3CR1^+^CD8^+^ T cells as an early on-treatment biomarker for clinical response to anti-PD-1 therapy.

## Results

### Increased frequency of PB CX3CR1^+^CD8^+^ T cells after effective ICI therapy

While ICI are known to restore cytokine production and proliferation of effector T cells^35^, whether or not ICI affect differentiation of tumor-specific T cells remains elusive. To this end, we evaluated CX3CR1 expression, a marker of T-cell differentiation in combination with CD27 (refs.^29–31^), on PB CD8^+^ T cells before and during treatment with anti-PD-L1 and anti-CTLA-4 Ab or isotype Ab (NT: no-treatment) in two mouse tumor models, MC38 and CT26 colon adenocarcinoma (**Fig. 1A**). Significant tumor growth delays (**Fig. 1B**) and increased frequency of CX3CR1^+^CD8^+^ T cells (**Fig. 1C**) were observed in MC38 and CT26 tumor-bearing mice treated with ICI compared to mice receiving isotype Ab. Next, we used a tetramer (Tet) to detect CD8^+^ T cells specific for mutated Adpgk protein (Adpgk^Mut^) in MC38 and shared tumor-associated antigen (TAA), gp70 in CT26 tumors^36, 37^, and found substantially increased frequency of CX3CR1^+^Tet^+^CD8^+^ T cells in both tumor models (**Fig. 1D**), suggesting that T-cell differentiation after ICI therapy occurs in tumor-specific CD8^+^ T cells.

**Figure 1.**
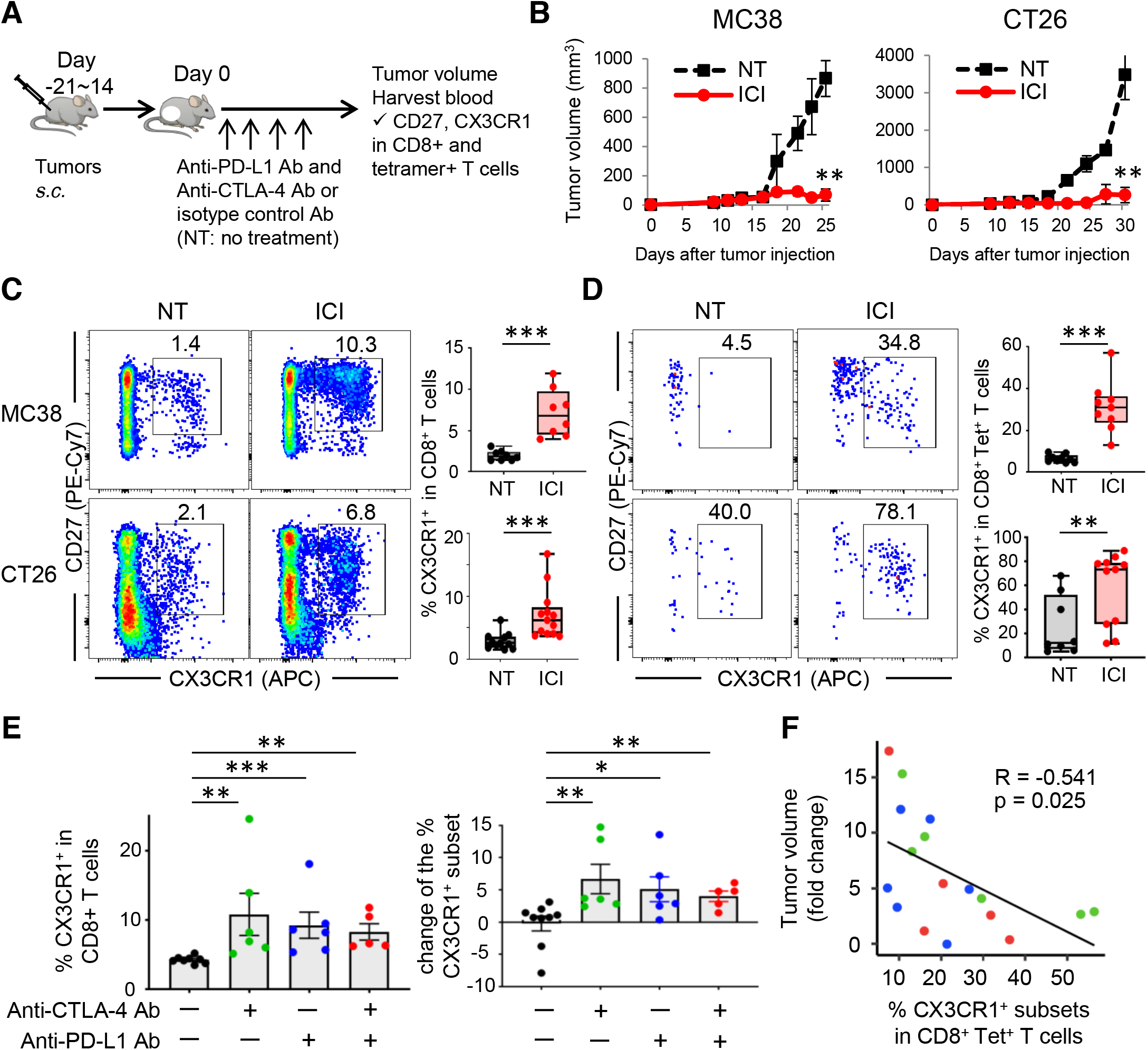
Changes in the frequency of PB CX3CR1^+^ CD8^+^ T cells associates with response to ICI therapy. **A,** Experimental scheme of treatment with immune checkpoint inhibitors (ICI). Anti-PD-L1 antibody (Ab) and anti-CTLA-4 Ab were administered intraperitoneally every 3 days and every other day, respectively. **B**, Tumor growth curves in MC38 (left) and CT26 (right) tumor-bearing mice treated with isotype Ab (NT) or ICI (n = 5 mice per group). **C, D**, Representative bivariate plot showing CD27 and CX3CR1 expression of CD8^+^ T cells (**C**) and tetramer (Tet)^+^ CD8^+^ T cells (**D**) in peripheral blood (PB) of MC38 (upper) and CT26 (lower) tumor-bearing mice in different treatments as indicated; numbers denote percent CX3CR1^+^ cells. Right panels show the frequency of CX3CR1^+^ cells among CD8^+^ T cells (**C**) and Tet^+^ CD8^+^ T cells (**D**). PB was harvested at day 13. n = 2 experiments pooled. **E**, Frequency of the CX3CR1^+^ subset among CD8^+^ T cells (left) and percent change of CX3CR1 ^+^ T-cell subsets (right) in PB of CT26 tumor-bearing mice treated with isotype control Ab (black), anti-CTLA4 Ab (green), anti-PD-L1 Ab(blue), or combination of these two Ab (red). (n = 5 - 7 per each group). PB was harvested at day 7. **F**, Scatter plot with correlation curve showing relationship between CT26 tumor volume (fold change) and the frequency of PB CX3CR1 ^+^Tet^+^CD8^+^ T cells at day 7 in CT26 tumor-bearing mice as in (E). Correlation is shown using Pearson correlation coefficients (R) and significance was determined using Spearman correlation. (**B – E**) *, *P* < 0.05; **, *P* < 0.01; ***, *P* < 0.001; Mann-Whitney *U*-test. Values are mean ± SEM.

### Changes in the frequency of PB CX3CR1^+^CD8^+^ T cells associate with response to ICI

Although both anti-PD-L1 Ab and anti-CTLA-4 Ab target subsets of exhausted-like CD8 T cells, they do so through distinct cellular mechanisms^38^. Therefore, we evaluated CX3CR1 expression of PB CD8^+^ T cells in CT26-bearing mice treated with either anti-PD-L1 Ab, anti-CTLA-4 Ab or both. Increased frequency as well as the percentage change of the PB CX3CR1^+^CD8^+^ T cells in individual mice were seen after either monotherapy or combined ICI therapy compared to no treatment (**Fig. 1E**). We also examined whether the frequency of the CX3CR1^+^ subset in Tet^+^CD8^+^ T cells correlates with treatment response. To this end, we drew PB and measured CT26 tumor size in individual mice before and 1 week after treatment with either anti-PD-L1 Ab, anti-CTLA-4 Ab or both. We found that frequency of the CX3CR1^+^ subset in PB Tet^+^CD8^+^ T cells correlated with response to ICI, suggesting the potential utility of CX3CR1 as a blood-based T-cell biomarker to predict response to ICI (**Fig. 1F**).

### CX3CR1 but not Ki-67 is stably upregulated in PB CD8^+^ T cells during ICI therapy

To gain insight into the PB CX3CR1^+^ CD8^+^ T cells, we evaluated the expression of the nuclear protein Ki-67, a marker for proliferation that is upregulated in subsets of PB CD8^+^ T cells in response to ICI^8, 9, 24–27^. The levels of Ki-67 expression in the CX3CR1^+^CD8^+^ T cells were significantly higher than in the CX3CR1^-^ (CD27^lo^CX3CR1^-^ and CD27^hi^CX3CR1^-^) subsets 2 weeks after ICI therapy in CT26 tumor-bearing mice (**Fig. 2A**). We next examined whether an increased expression of CX3CR1 on PB CD8^+^ T cells is transient or sustained during ICI therapy. Increased frequency of CX3CR1^+^ subset was seen in both CD8^+^ and Tet^+^CD8^+^ T cells starting from day 7, which remained high during treatment (**Fig. 2B**). In contrast, Ki-67 expression peaked at day 14, and returned to the baseline at day 21 (**Fig. 2B**).

**Figure 2.**
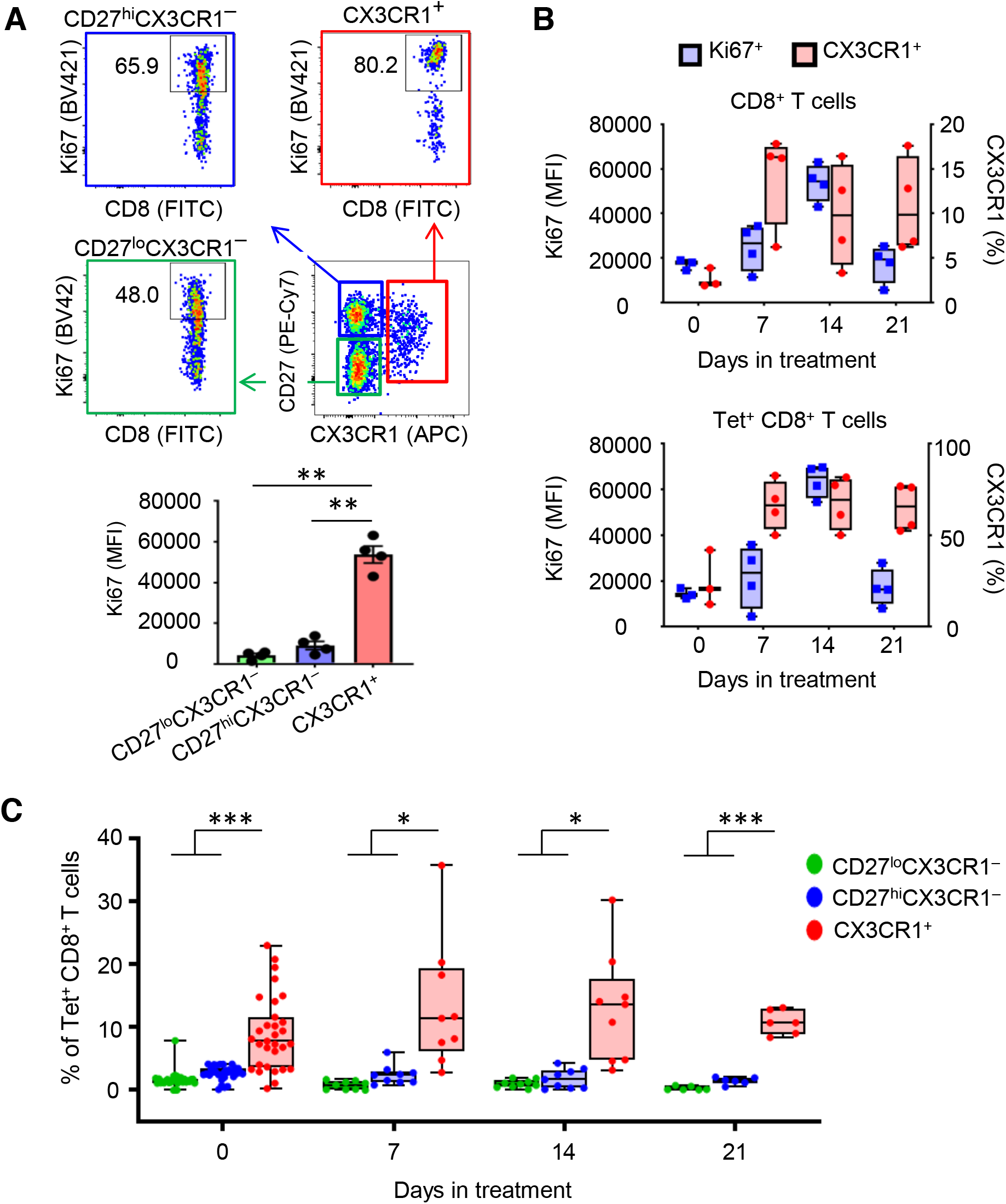
Phenotypic analysis of PB CX3CR1^+^ CD8^+^ T cells before and during ICI therapy. **A-C,** CT26 tumor-bearing mice were treated with anti-CTLA-4 Ab and anti-PD-L1 Ab. Peripheral blood (PB) was obtained before and during treatment. **A**, Ki-67 expression of CD27^lo^CX3CR1^-^ (green), CD27^hi^CX3CR1^-^ (blue), and CX3CR1^+^ (red) CD8^+^ T cells in PB 2 weeks after ICI therapy. Numbers denote percent Ki-67^+^ cells. Data panel shows mean fluorescence intensity (MFI) of Ki-67^+^ cells in each subset (n=4 per each group). **B**, Box and whiskers plots showing MFI of Ki-67^+^ (blue) and frequency of the CX3CR1^+^ subset (red) in CD8^+^ T cells (upper) and Tet^+^CD8^+^ T cells (lower) at day 0, 7, 14, and 21 in PB (n = 3 - 4 per each group). **C**, Frequency of Tet^+^CD8^+^ T cells in the CD27^lo^CX3CR1^-^ (green), CD27^hi^CX3CR1^-^ (blue), and CX3CR1^+^(red) subsets at day 0, 7, 14, and 21 in PB. n = 2 experiments pooled. (**A** and **C**) *, *P* < 0.05; **, *P* < 0.01; ***, *P* < 0.0005; repeated-measures one-way ANOVA with Tukey’s multiple comparisons test. Values are mean ± SEM.

### Tumor-specific CD8^+^ T cells are enriched in the CX3CR1^+^ subset in PB

Identification of circulating T-cell biomarkers that enrich tumor-reactive T cells may facilitate discovery of dynamic predictive marker of response to ICI. First, we assessed the frequency of Tet^+^CD8^+^ T cells within the PB CX3CR1^+^ and CX3CR1^-^ subsets in MC38 and CT26 tumor-bearing mice (**Supplementary Fig. 1A**). We found more Tet^+^CD8^+^ T cells in the CX3CR1^+^ subset than in the CX3CR1^-^ subsets even before treatment although the frequency varied between individual mice (**Supplementary Fig. 1B**). Next, we evaluated change in the frequency of Tet^+^CD8^+^ T cells during ICI therapy in CT26 tumor-bearing mice. The frequency of Tet^+^CD8^+^ T cells remained higher in the CX3CR1^+^ subset than in the CX3CR1^-^ subsets, and became less variable between individual mice at day 21 (**Fig. 2C**).

### Clonally expanded TCR repertoires of CD8^+^ TILs are enriched in the peripheral CX3CR1^+^ subset during ICI therapy

The high frequency of tumor-specific CD8^+^ T cells in the CX3CR1^+^ subset before and during ICI treatment is suggestive that heterogeneous tumor-infiltrating CD8^+^ T cells are also enriched in the CX3CR1^+^ subset. To this end, we performed TCR sequencing on isolated CD8^+^ TILs and splenic CD27^lo^CX3CR1^-^, CD27^hi^CX3CR1^-^, and CX3CR1^+^CD8^+^ T cells from MC38 tumor-bearing mice treated with combined ICI therapy (**Supplementary Fig. S2A, B**). Comparison of the TCR repertoire in CD8^+^ TILs and three subsets of splenic CD8^+^ T cells demonstrated high degree of overlap in TCR usage between CD8^+^ TILs and splenic CX3CR1^+^CD8^+^ T cells (**Fig. 3A; Supplementary Fig. S3**) as determined by the Morisita’s overlap index^39^.

**Figure 3.**
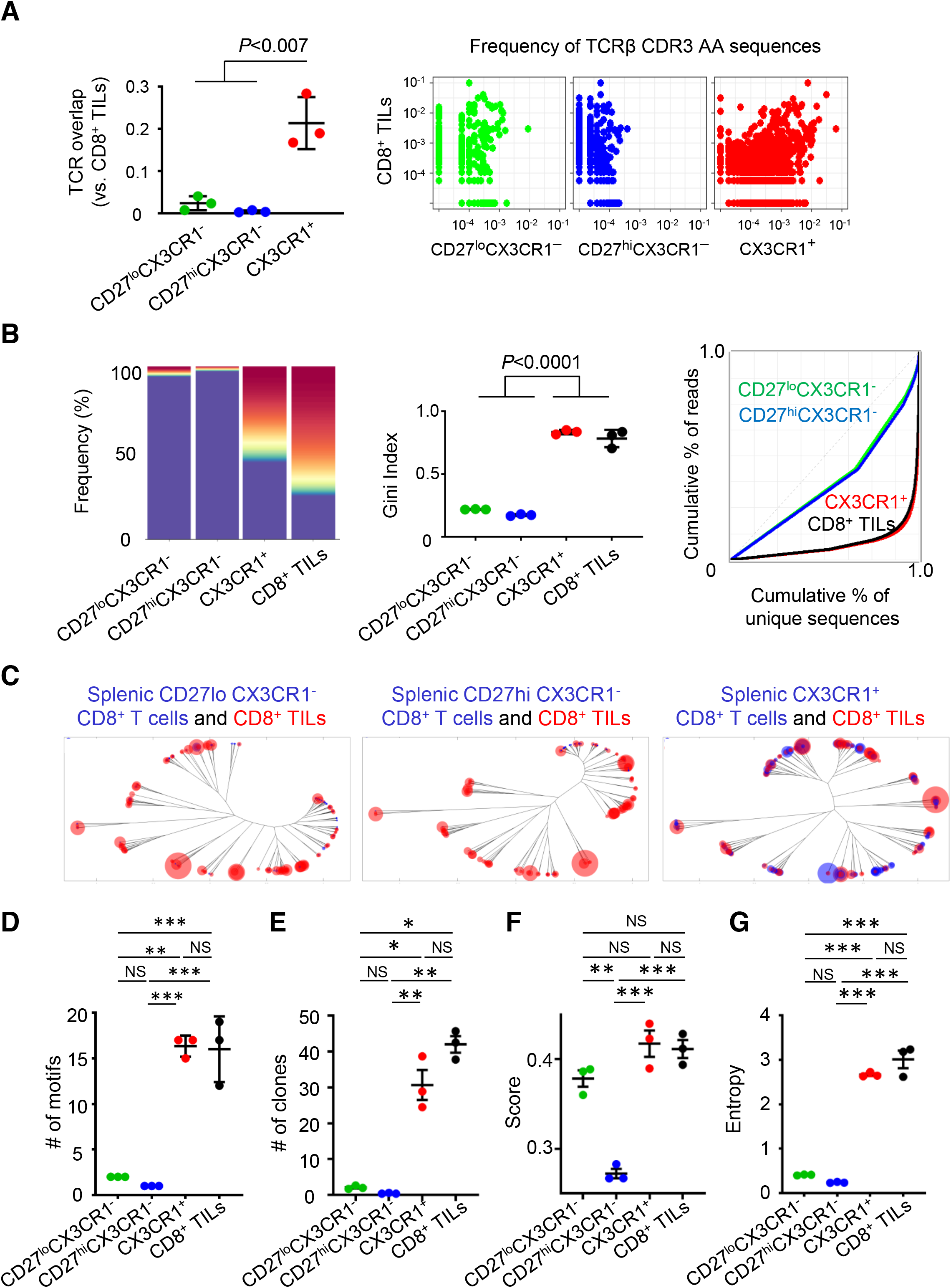
Effective ICI therapy induces high degree of TCR sequence similarity and clonality between tumor infiltrating CD8^+^ T cells and peripheral CX3XR1 ^+^ CD8^+^ T cells. **A-G,** MC38 tumor-bearing mice were treated with anti-CTLA-4 Ab and anti-PD-L1 Ab. Three subsets of splenic CD8^+^ T cells determined by CD27 and CX3CR1 expression (CD27^lo^CX3CR1^-^, CD27^hi^CX3CR1^-^, and CX3CR1^+^), and CD8^+^ tumor-infiltrating lymphocytes (TILs) were isolated 2 weeks after treatment for TCR repertoire and clonality analysis. **A**, TCR repertoire overlap by Morisita’s index (left) and pairwise scatter plots of the frequency of TCRβ CDR3 amino acid (AA) sequences between each subset of splenic CD8^+^ T cells and CD8^+^ TILs (right). **B**, TCR clonality analysis of three subsets of splenic CD8^+^ T cells and CD8^+^ TILs by top sequence plot (left), Gini index (center), and Lorenz curve (right). The most abundant 100 AA sequences are colored while other less frequent clones are in purple in top sequence plot. (**A** and **B**) Data were analyzed using the one-way ANOVA test with Tukey’s multiple comparisons to generate *P* values. **C**, Representative overlapped weighted TCR repertoire dendrograms by ImmunoMap analysis between three subsets of splenic CD8^+^ T cells (blue) and CD8^+^ TILs (red). The distance of the branch ends represents sequence distance, and the size of circles denotes frequency of sequence. Data shown are representative of three independent experiments. **D-G**, Number of dominant motifs (**D**), number of clones contributing response (**E**), TCR diversity score (**F**) and Shannon’s entropy (**G**) of splenic CD27^lo^ CX3CR1^-^ (green), CD27^hi^ CX3CR1^-^ (blue), CX3CR1^+^ (red) CD8^+^ T cells, and CD8^+^ TILs (black). NS, not significant, *, *P* < 0.05; **, *P* < 0.005; ***, *P* < 0.0005 by repeated-measures one-way ANOVA with Tukey’s multiple comparisons test. Values are mean ± SEM.

Next, we evaluated TCR clonality in CD8^+^ TILs and three subsets of splenic CD8^+^ T cells 2 weeks after ICI therapy (**Fig. 3B**). The 100 most abundant TCR clones (colored) comprised more than 50% of TCR sequences in the splenic CX3CR1^+^ subset and CD8^+^ TILs while the majority of TCR sequences in the splenic CX3CR1^-^ subsets were constituted of less frequent clones (purple). To quantify the skewness of the clonal distribution, we measured the Gini index, and found similarly higher clonality in the splenic CX3CR1^+^ subset and CD8^+^ TILs compared to splenic CX3CR1^-^ subsets. Lorenz curves for splenic CX3CR1^+^ subset and CD8^+^ TILs were far from the equidistribution line, suggesting unequal distribution and skewing of the TCR repertoire.

### ICI therapy induces high degree of TCR sequence similarity and clonality between CD8^+^ TILs and the peripheral CX3CR1^+^ subset

TCRs that recognize the same antigen may not be the exact TCR clonotypes but have highly homologous sequences and share similar sequence features^40, 41^. Unlike Morisita’s overlap index, a bioinformatics program, ImmunoMap^42^, allows us to analyze biological sequence similarity in between peripheral and intratumoral CD8^+^ T cells. Compared to the CX3CR1^-^ subsets, splenic CX3CR1^+^CD8^+^ T-cell subsets contain highly frequent clonally expanded CD8^+^ T cells indicated by the large circles in the end of many different branches of the tree 2 weeks after ICI therapy (**Fig. 3C**; **Supplementary Fig. S4A, B**).

Analysis of the top six dominant CDR3β amino acid (AA) sequences in splenic CX3CR1^+^CD8^+^ T cells and CD8^+^ TILs revealed they shared a highly frequent AA sequence (CASSLVGNQDTQYF) in all three independent experiments (**Table 1, yellow highlighted**). Although the most frequent AA sequence, <u>CASSP</u>RLGDN<u>YAEQFF</u>, in splenic CX3CR1^+^CD8^+^ T cells was not identified in CD8^+^ TILs in the same experiment (Exp. #1 in **Table 1A**), this AA sequence had a high degree of sequence homology with dominant AA sequences, <u>CASSP</u>G<u>YAEQFF</u> and <u>CASSP</u>GQG<u>YAEQFF</u> in CD8^+^ TILs, located in the same branch of the dendrogram (**Table 1B, blue highlighted; Supplementary Fig. S4A**). Similarly, abundant AA sequences, <u>CASS</u>PGRG<u>YEQYF</u> in splenic CX3CR1^+^CD8^+^ T cells and <u>CASSS</u>GT<u>YEQYF</u> in CD8^+^ TILs clustered largely in the same branch, indicating that they shared a high degree of similarity (Exp. #2 in **Table 1, green highlighted; Supplementary Fig. S4B**).

**Table 1:**
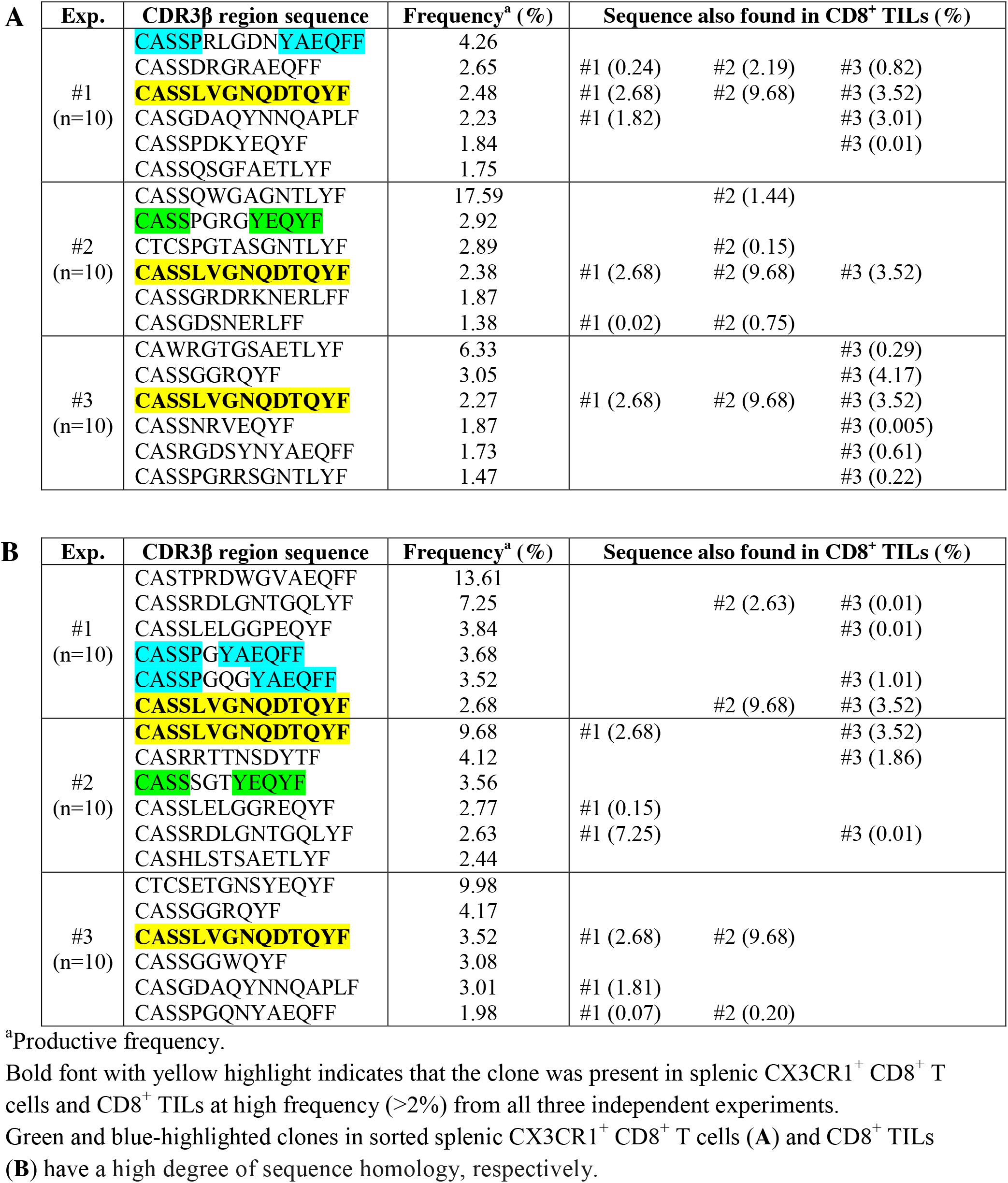
Six most dominant clones in sorted splenic CX3CR1^+^ CD8^+^ T cells (A) and CD8^+^ TILs (B) in MC38-bearing mice treated with CTLA-4 and PD-L1 blockades (n=10 mice / experiment)

Furthermore, CD8^+^ TILs and splenic CX3CR1^+^CD8^+^ T cells had a similar number of structural motifs that dominated the response, and higher numbers of responding clones that expanded 10 times more than the summation of all other homologous clones in a sample (**Fig. 3D, E**). The TCR diversity score and the Shannon entropy calculations revealed that both populations have similar TCR diversity (**Fig. 3F, G**). Collectively, these findings suggest TCR repertoires in peripheral CX3CR1^+^CD8^+^ T-cell clones reflect the TCR repertoires in CD8^+^ TILs, and CX3CR1 on PB CD8^+^ T cells may act as a dynamic biomarker during the course of effective ICI therapy.

### Expansion of the CX3CR1^+^ subset in PB CD8^+^ T cells correlates with improved response to anti-PD-1 therapy and survival in patients with NSCLC

We next explored whether changes in the frequency of the CX3CR1^+^ subset in PB CD8^+^ T cells correlate with response to ICI in patients. We analyzed PBMC samples from the PB of a cohort of 36 NSCLC patients treated with anti-PD-1 Ab (Pembrolizumab or Nivolumab) (**Fig. 4A**). PB was obtained before treatment and every 2-6 weeks during therapy for 12 weeks. All patients had pretreatment tumor tissue available to assess PD-L1 expression. The frequency of TILs and TMB were also analyzed as described before^43^. Baseline characteristics of 36 NSCLC patients are described in **Supplementary Table S1**. Clinical response in individual patients was derived from investigator-reported data per iRECIST criteria^44^ at the 12 week time point. Overall response rates (ORR) which include complete response (CR) and partial response (PR) were 36.7% and 20% for patients with a PD-L1 tumor proportion score (TPS) of 50% or greater and 1-49%, respectively, in line with previous studies^45, 46^.

**Figure 4.**
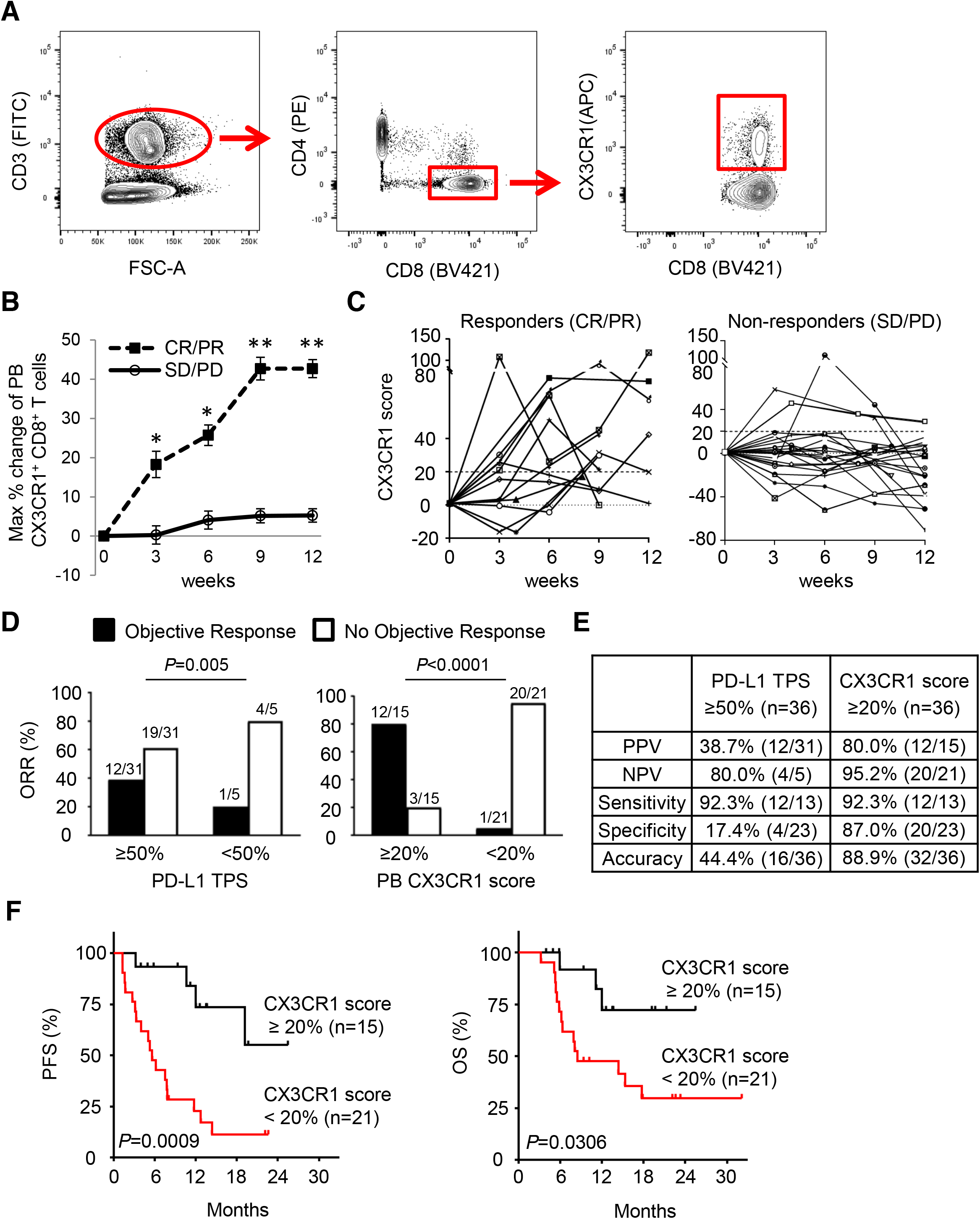
Expansion of the CX3CR1^+^ subset in PB CD8^+^ T cells correlates with improved response to anti-PD-1 therapy and survival in patients with NSCLC. **A**, Gating strategy for identifying CX3CR1 ^+^ CD8^+^ T cells in peripheral mononuclear blood cells. Cells were first gated for lymphocytes (SSC-A vs. FSC-A) and for singlets (FSC-H vs. FSC-A). **B**, Maximal % change of CX3CR1^+^ subset in PB CD8^+^ T cells in responders and non-responders of 36 NSCLC patients treated with anti-PD-1 therapy by 12 weeks. CR/PR: complete and partial response, SD/PD: stable and progressive disease. * *P*<0.04, \*\**P*<0.0001 by Mann-Whitney *U*-test. Values are median ± SEM. **C**, Percent change of the CX3CR1 ^+^ subset in PB CD8^+^ T cells from baseline (CX3CR1 score) in responders (left) and non-responders (right). **D**, Objective response rate (ORR) for high and low PD-L1 tumor proportion score (TPS) (left) and PB CX3CR1 score (right). ORR was analyzed by Fisher’s exact test. **E**, Comparison of biomarker performance between PD-L1 TPS and CX3CR1 score at 12 weeks. **F**, Kaplan-Meier progression free survival (PFS) (left) and overall survival (OS) (right) for high versus low CX3CR1 score. The two-tailed *P* value was calculated using the log-rank test.

We evaluated the maximal percent change of the CX3CR1^+^ subset in PB CD8^+^ T cells at 3, 6, 9 and 12-weeks relative to baseline. The maximal percent change was substantially higher in responders than non-responders as early as 3 weeks from the initiation of the treatment (**Fig. 4B**). Next, we obtained estimates of the area under the curve (AUC) and corresponding 95% confidence interval (CI) using a logistic regression model, and the optimal cut-off score for discriminating between groups using the Youden’s index criterion^47^. These analyses revealed that an increase of CX3CR1^+^CD8^+^ T-cell subsets by 15.5~21.2% from baseline segregated responders from non-responders at 6-12 weeks (**Supplementary Table S2**). Hereafter the percent change of the CX3CR1^+^ subset in PB CD8^+^ T cells from baseline are designated as a “CX3CR1 score”. **Fig. 4C** shows longitudinal CD8 T-cell responses in individual patients. We found at least 20% increase of the CX3CR1 score in 92.3% (12/13) of responders compared to only 13.0% (3/23) of non-responders.

Based on these results, we hypothesized that a cut-off of at least 20% increase of the CX3CR1 score would correlate with response to anti-PD-1 therapy and assessed the association between the CX3CR1 score and objective response. The maximal CX3CR1 score of at least 20 was strongly associated with ORR (*P*<0.0001) (odds ratio, 80.00; 95% CI, 7.45 to 858.94), while PD-L1 TPS at ≥50% was also correlated with ORR (*P*=0.005) (odds ratio, 2.53; 95% CI, 0.25 to 25.39) (**Fig. 4D**). Next, we analyzed the corresponding sensitivity, specificity, positive predictive value (PPV), and negative predictive value (NPV) of the CX3CR1 score and the PD-L1 TPS. The CX3CR1 score demonstrated remarkably high PPV, NPV, sensitivity, and specificity, and identified response in 32/36 (88.9%) while a PD-L1 TPS of at least 50% had suboptimal PPV and specificity, and correctly identified response only in 16/36 (44.4%) (**Fig. 4E**). Notably, tumor tissues were available for assessing frequency of TILs and TMB for only 66.6% (24/36) and 61.1% (22/36) of NSCLC patients, respectively (**Supplementary Table S3**), suggesting limitation of these analyses, in line with previous reports^48, 49^. Lastly, we evaluated correlation between the CX3CR1 score and survival. We found that at least 20% increase of the CX3CR1 score was associated with better progression free (PFS) and overall survival (OS) (**Fig. 4F**). Median PFS and OS among patients with a CX3CR1 score of less than 20% were 5.7 months (95% CI, 2.2 to 11.8), and 8.6 months (95% CI, 1.4 to 9.4), respectively, while median PFS and OS among patients with a CX3CR1 score of at least 20% were not reached. Taken together, the CX3CR1 score provides highly accurate prediction of a patient’s clinical response early on-treatment and correlates with survival.

## Discussion

The lack of robust predictive biomarker for response is a major obstacle of ICI therapy. Blood-based mechanism-driven dynamic biomarkers that reflect constantly evolving TME would be ideal, and intense efforts are ongoing to identify circulating biomarkers for ICI response^4–7, 19–23^. Here, we provide evidence in tumor-bearing mice that: 1) effective ICI therapy correlates with the increased frequency and TCR clonality of peripheral CX3CR1^+^CD8^+^ T cells that identify an enriched repertoire of neoantigen- and TAA-specific CD8^+^ T cells; 2) the frequency of CX3CR1^+^ but not Ki-67^+^ PB CD8^+^ T cells remained elevated during ICI therapy; and 3) there are high degree of TCR sequence overlap and similarity between CD8^+^ TILs and the peripheral CX3CR1^+^ subset during ICI therapy. Furthermore, analysis of longitudinal PB samples obtained from a cohort of NSCLC patients highlights the potential clinical utility of CX3CR1 as a useful blood-based biomarker to predict response to ICI early after initiation of therapy.

Mechanistically, there are some potential advantages for CX3CR1 as a blood-based biomarker. First, unlike other molecules such as Ki-67 and PD-1 which are transiently upregulated on T cells after activation, CX3CR1 is stably expressed on virus- and tumor-specific CD8^+^ T cells upon differentiation^30–32^. In agreement with this, we found CX3CR1 but not Ki-67 remains elevated in tumor-specific CD8^+^ T cells during ICI therapy in preclinical models. Additionally, many responding NSCLC patients in our cohort maintained the level of CX3CR1 expression on CD8^+^ T cells above their baseline. Second, given the low levels of CXCR3 expression in CX3CR1^hi^ CD8^+^ T cells^30^, which is required to traffic to tumors^50^, it is possible that CX3CR1^+^CD8^+^ T cells remain in circulation, and might not actively traffic to the tumor unless fractalkine (CX3CL1), the ligand of CX3CR1 is produced from the TME. In support with this notion, a recent study showed a higher fraction of CX3CR1^+^CD8^+^ T cells in PB compared with tumors in NSCLC patients^51^. Furthermore, we have recently reported that adoptively transferred tumor-specific CX3CR1^-^CD8^+^ T cells generate CX3CR1^+^CD8^+^ T cells upon *in vivo* stimulation, traffic to the TME, and mediate effective regression of established tumors^32^. In contrast, CX3CR1^+^CD8^+^ T cells had no impact on established tumors compared with the CX3CR1^-^ subset and became dominant in PB in a preclinical model^32^. These features of CX3CR1^+^CD8^+^T cells might have contributed to the greater accuracy of the CX3CR1 score in our cohort and make them uniquely suitable for a circulating T-cell biomarker.

Increased TCR clonality can be identified in tumors and associates with response to anti-PD-1 therapy^10^; however, the role of PB TCR clonality in predicting response to ICI remains elusive. PB T cells contain highly diverse TCR repertoires, the majority of which are not specific to the tumor; therefore, changes of PB TCR clonality might be difficult to detect even in responders. Identification of markers to detect tumor-specific T cells in PB might overcome this limitation. PD-1 was found to be a cell-surface marker to identify PB neoantigen-reactive T cells^52^, and a recent study showed TCR clonality changes can be observed in PB PD-1^+^CD8^+^ T cells^53^. Although additional clinical studies are required to determine the predictive and prognostic significance of the TCR clonality in PB CX3CR1^+^CD8^+^ T cells, our findings also provide insight into the utility of TCR clonality in the peripheral T-cell subset. Our findings of high clonality in peripheral CX3CR1^+^CD8^+^ T cells that contain intratumoral CD8^+^ T-cell repertoires also align with emerging evidence from high-dimensional profiling of PB T cells by single-cell RNA and/or TCR sequencing showing expansion of cytotoxic effector memory CD8^+^ T cells that include novel intratumoral clones in patients responding to ICI therapy^22, 54, 55^.

There is an unmet clinical need in establishing reliable predictive biomarkers for combined anti-PD-1 and anti-CTLA-4 therapy, where the risk of severe immune-related adverse events (irAEs) is as high as the proportion of patients responding to combined ICI. High NPV early on-treatment biomarkers could reliably identify a lack of response, and minimize unnecessary treatment, toxicity and associated cost. This is especially relevant for patients with NSCLC who have various treatment options available. Our analyses demonstrated that the CX3CR1 score ≥20 at 6-12 weeks could discriminate responders from non-responders with high NPV, suggesting that the CX3CR1 score may help make a decision about whether to continue ICI or switch to alternative more effective treatment.

The CX3CR1 score might be convenient to use in clinical setting from technical and analytical perspective. While we isolated PBMCs in this study, the same analysis can be done with as little as 1 ml of whole blood without isolating PBMC by density gradient centrifugation. Unlike intracellular proteins, there is no need for fixation/permeabilization procedure to stain CX3CR1. The CX3CR1^+^ subset can be easily distinguishable from the CX3CR1^-^ subsets in PB CD8^+^ T cells unlike other markers such as PD-1, where the boundary between positive and negative populations might be difficult to set^56^. The CX3CR1 score can be obtained by traditional fluorescence flow cytometric analysis, and does not require NGS platform, complex algorithm, or bioinformatics analysis. Thus, the results can be readily available for prompt feedback to oncologists and patients.

Tumor PD-L1 expression predicts the likelihood of response to ICI and is used as a companion diagnostic for NSCLC but not for several other malignancies such as melanoma. Therefore, it remains elusive whether the CX3CR1 score would be useful in patients with other types of cancer. A recent study showed T-cell invigoration to tumor burden ratio was associated with response to anti-PD-1 Ab and clinical outcome in melanoma patients^24^. Although it was not within the scope of our study, one future area of investigation would be to compare the utility of different PB T-cell biomarkers in the same disease, or evaluate whether combining multiple biomarkers could improve predictive and prognostic values.

In summary, our findings demonstrate that CX3CR1 is a useful dynamic circulating T-cell biomarker, and that at least 20% increase of the CX3CR1^+^ subset in PB CD8^+^ T cells identifies NSCLC patients responding to anti-PD-1 therapy early on-treatment. Our study provides a rationale for further investigation to test the utility of the circulating T-cell differentiation marker for a wide variety of malignancies in larger prospective trials.

## Methods

Please see the Supplementary Data.

## Acknowledgments

We thank the patients, their families and the clinical study staff, Ms. Elongia Farrell, Noelle Brunsing, Suzanne M. Stack, Rushka Kallicharan-Smith, and Ann Marie DiRaddo (Roswell Park) for their contribution to the study. We acknowledge the NIH Tetramer Core Facility (contract HHSN272201300006C) for provision of MHC-I tetramers, and Dr. Weiping Zou (University of Michigan, Ann Arbor, Michigan) for MC38 cells, Drs. James J. Moon (University of Michigan, Ann Arbor, Michigan), Kenichi Makino and Toshihiro Yokoi (Roswell Park) for technical assistance, and Ms. Sue Hess and Judith Epstein (Roswell Park) for administrative assistance. This work was supported by National Cancer Institute (NCI) grant P30CA016056 involving the use of Roswell Park’s Flow and Image Cytometry, Clinical Research and Pathology Network, Biostatistics & Statistical Genomics Shared Resource and Genomic Shared Resources, and by the National Center for Advancing Translational Sciences of the NIH (UL1TR001412) to the University of Buffalo. This work was supported by Roswell Park Alliance Foundation, the Melanoma Research Alliance, Department of Defense Lung Cancer Research Program (LC180245), and NCI grant, K08CA197966 (F. Ito), R01CA188900 (B.H. Segal), R01CA188900 (B.H. Segal), U01CA154967, P50CA159981-01A1 (K. Odunsi), Uehara Memorial Foundation (T. Oba), and Astellas Foundation for Research on Metabolic Disorders and the Nakatomi Foundation (T. Yamauchi).

## Supplementary Materials and Methods

### Mice

Male and female C57BL/6 mice and female Balb/c mice were purchased from the Jackson Laboratories. All mice were 7 to 12 weeks old at the beginning of each experiment, and were housed in the Unit for Laboratory Animal Medicine at the Roswell Park Comprehensive Cancer Center in compliance with the Institutional Animal Care and Use Committee regulations.

### Cell lines

MC38 and CT26 murine colon adenocarcinoma cell lines were gifts from Dr. Weiping Zou (University of Michigan) and Dr. Sharon Evans (Roswell Park Comprehensive Cancer Center), respectively. MC38 and CT26 cells were cultured in RPMI (Gibco) supplemented with 10% FBS (Sigma), 1% NEAA (Gibco), 2 mM GlutaMAX-1 (Gibco), 100 U/ml penicillin-streptomycin (Gibco), and 55 μM 2-mercaptoethanol (Gibco). Cells were authenticated by morphology, phenotype and growth, and routinely screened for *Mycoplasma*, and were maintained at 37°C in a humidified 5% CO_2_ atmosphere.

### *In vivo* mouse studies

Male or female C57BL/6 mice and female Balb/c mice were inoculated with 5-8 × 10^5^ MC38 and 5 × 10^5^ CT26, respectively per mouse on the right flank by subcutaneous injection on day 0. When tumor volume reached approximately 50 mm^3^, 200 μg of anti-PD-L1 Ab (clone 10F.9G2, BioXcell) and/or 100 μg of anti-CTLA-4 Ab (clone 9H10, BioXcell) were administered intraperitoneally every 3 days and every other day, respectively. Polyclonal syrian hamster IgG (BioXcell) and rat IgG2b, κ (BioXcell) were used as isotype control Abs. Tumor volumes were calculated by determining the length of short (*l*) and long (*L*) diameters (volume = l^2^ x L/2). Experimental end points were reached when tumors exceeded 20 mm in diameter or when mice became moribund and showed signs of lateral recumbency, cachexia, lack of response to noxious stimuli, or observable weight loss.

### Single-cell preparations

Blood, spleens and tumors were harvested at day 14-42 post MC38 or CT26 tumor implantation. Spleens were homogenized by forcing the tissue through a cell strainer (70 μm; BD Biosciences). Red blood cells in blood and spleen were lysed using ACK Lysis Buffer (Gibco). Tumors were cut into small pieces of 2-4 mm. Single-cell suspensions were obtained by mechanical dispersion consisting of two 30-min incubations at 37°C, 5% CO_2_ in 5 ml RPMI 1640 (Gibco) and tumor dissociation kit (Miltenyi Biotec) in C Tubes (Miltenyi Biotec) interspersed with three mechanical dispersions on a GentleMACS dissociator (Miltenyi Biotec). The tumor cell suspensions were then filtered through a cell strainer (70 μm; BD Biosciences).

### Flow cytometry and cell sorting

Surface staining of leukocytes in murine blood, spleens and tumors was performed in FACS buffer (made in house) using monoclonal antibodies against mouse CD3 (145-2C11), CD90.2 (53-2.1), CD4 (GK1.5), CD8 (53-6.7), CX3CR1 (SA011F11) (all BioLegend), CD27 (LG.7F9, eBioscience), CD45 (30-F11, Invitrogen), CD8 (KT15 for tetramer staining, Invitrogen), and PD-1 (J43, BD Biosciences). Live/dead cell discrimination was performed using Live/Dead Fixable Aqua Dead Cell Stain Kit or LIVE/DEAD Fixable Near-IR Dead Cell Stain Kit (Invitrogen). Samples were incubated with antibodies for 20 min at RT in the dark. We used the tetramer staining assay with peptide-MHC tetramer tagged with PE (H-2D^b^-restricted ASMTNMELM for MC38-bearing mice and H-2Ld-restricted SPSYVYHQF for CT26-bearing mice (The NIH Tetramer Core Facility)) to analyze the percentages of tumor antigen-specific CD8^+^ T cells. For intracellular staining, surface-stained cells were fixed and permeabilized using a Foxp3 fixation/permeabilization kit (eBioscience), then stained with anti-Ki67 (16A8, BioLegend) for 30 min.

For TCR sequencing of murine splenic CD8^+^ T cells, single cell suspensions from freshly isolated splenocytes were stained as above. CD45^+^ CD3^+^ CD8^+^ T cells were gated, and CD27^lo^ CX3CR1^-^, CD27^hi^ CX3CR1^-^, and CX3CR1^+^ CD8^+^ T cells were sorted using BD Aria Sorter. An EasySep Mouse CD8a Positive Selection Kit II (STEMCELL Technologies) was used to isolate murine CD8^+^ TILs for TCR sequencing.

For phenotypic analysis of PBTCs, fresh or cryopreserved PBMC samples were stained with master mix of antibodies for surface stains including CD3 (UCHT1, BD Biosciences), CD4 (SK3, BD Biosciences), CD8 (RPA-T8, eBioscience), CD27 (O323, eBioscience), and, CX3CR1 (2A9-1, Biolegend). Samples were acquired using LSR II (BD), LSRFortessa (BD) or SONY sorter and data analyzed with FlowJo software v10.1.5 (TreeStar).

### DNA isolation, TCRβ CDR3 region sequencing and repertoire analysis

DNA from flow-isolated murine splenic CD8^+^ T cells and CD8^+^ TILs, and PB CD8^+^ T cells was extracted using (QIAamp DNA Micro Kit (QIAGEN)). DNA was quantified using Qubit dsDNA BR Assay (Invitrogen). Amplification and sequencing of TCRβ CDR3 regions was performed using ImmunoSEQ immune profiling system at the survey level (Adaptive Biotechnologies)(1). Sequencing was performed on an Illumina NextSeq system using 150 cycle mid-output kit (Illumina Inc.). Processed data were uploaded to the immunoSEQ Platform (Adaptive Biotechnologies) for preliminary bioinformatics analysis. Processed data were downloaded and frequencies/counts for TCR clonotypes and diversity were examined by nucleotide sequences after non-productive reads were filtered out.

TCR beta chain CDR3 variable region sequencing was performed using the ImmunoSEQ assay at the survey level (Adaptive Biotechnologies). T-cell repertoires, comprising all detected CDR3 sequences with annotated V and J gene segment identifications were downloaded directly from the ImmunoSEQ Analyzer from Adaptive biotechnologies. Metrics of the complete TCR repertoire in each sample, including the number of productive rearrangements, productive clonality and clonal frequencies were determined using the ImmunoSEQ Analyzer software and confirmed using the LymphoSeq package (2). All other analyses were performed using the LymphoSeq package and custom scripts in the R statistical software environment. Dissimilarity between sample repertoires was calculated using the Morisita’s Index(3), using the vegan package. Differential clone frequencies between samples were determined using the Fisher’s exact test with multiple test correction (Holm method). For differential analysis, only those clones observed with at least 5 cumulative read counts were considered. TCR clonality was calculated as 1-Pielou’s evenness(4) using the immunoSEQ Analyzer®. Clonality values approaching 1 indicate a very skewed distribution of frequencies, whereas values approaching 0 indicate that every rearrangement is present at nearly identical frequency.

TCR repertoires were visualized as weighted dendrograms using ImmunoMap(5). Only productive sequences with a frequency > 0.1% in the tumor were considered for analysis. Sequence distances were calculated based on sequence alignments scores using a PAM10 scoring matrix and gap penalty of 30. Circles are overlaid at the end of the branches corresponding to the CDR3 sequences with diameters proportional to the frequency of the sequences observed in the samples. Shannon’s Entropy, Dominant motifs, singular structural clones, singular clones contributing response, and richness of motifs were identified using ImmunoMap.

### Data reporting

No statistical methods were used to predetermine sample size. The experiments were not randomized and the investigators were not blinded to allocation during experiments and outcome assessment.

### Study design, patients and specimen collection

Thirty-six patients with naive or previously treated PD-L1 IHC positive non-small cell lung cancer (NSCLC) adenocarcinoma and squamous cell type, undergoing anti-PD-1 Ab (Pembrolizumab or Nivolumab) (Supplemental Table S2) were consented to the collection and storage of blood samples, the analysis of archived tumor tissue, and the review of their medical records under the protocol (I 188310), in accordance with the Institutional Review Board of Roswell Park Comprehensive Cancer Center. Peripheral blood was obtained in EDTA-containing tubes before treatment and before each infusion and every 2-6 weeks for 12 weeks. Peripheral blood mononuclear cells (PBMCs) were isolated using Lymphocyte Separation Medium (Corning) density gradient centrifugation and stored using standard protocols.

### Assessment of response

Clinical response to anti-PD-1 therapy was determined as best response based on immune related RECIST (iRECIST)(6) at the 12 week time point, and classified as complete response (CR) and partial response (PR) for responders or stable disease (SD) and progressive disease (PD) for non-responders. Objective responses were confirmed by at least one sequential tumor assessment, and objective response rates were calculated as [(CR + PR) ÷ number of patients] × 100. Fisher’s exact test was used to assess the association between PD-L1 expression and objective response.

### Immunohistochemical studies

The expression of PD-L1 on the surface of tumor cells and frequency of CD8^+^ T cells were evaluated as described before (7). Briefly, the expression of PD-L1 on the surface of tumor cells was assessed by means of the Dako Omnis platform (Agilent) with the 28–8 pharmDx antibody and scored by published guidelines (8). Serially sectioned tissue was evaluated for lymphocyte infiltration using the anti-CD8 antibody C8/144B (Agilent) and assigned a qualitative score of non-infiltrated, infiltrated, or excluded. Non-infiltrated referred to a sparse number of CD8^+^ T-cells that infiltrate nests of neoplastic cells and with less than 5% of the tumor showing an infiltrating pattern. Infiltrated represents frequent CD8^+^ T-cells that infiltrate nests of neoplastic cells in an overlapping fashion at least focally and in more than 5% of the tumor. Excluded represents restriction of more than 95% of all CD8^+^ T-cells in a tumor to the periphery or interstitial stromal areas and not actively invading nest or groups of neoplastic cells.

### Tumor mutational burden profiling

Tumor mutational burden (TMB) was evaluated as described before (7). In brief, DNA was extracted from each sample and processed for whole-exon DNAseq. TMB was assessed by targeted capture and sequencing of 409 cancer-related genes and amplicon sequencing of 394 immune transcripts, respectively, comprising 1.4 Mb of DNA on samples that met validated quality control (QC) thresholds (7). Somatic mutation calling was conducted using Ion Torrent Suite software’s variant caller plugin. Mutational burden (MuB) cutoff was derived from a reference population whereby the median MuB was determined on regular basis. This value was used as a baseline and a high MuB was defined as 2× this median value, or a value of 10.0.

### Statistics

Statistical analysis was performed using *t*-test or Mann-Whitney *U* test for comparisons between 2 groups, 1-way repeated measures ANOVA with Turkey-adjusted multiple comparisons for comparisons more than 2 groups, or the Mantel-Cox method (log-rank test) for survival analysis using GraphPad Prism 7.03 (GraphPad Software) and the R statistical software. TCR repertoire analysis was performed using ImmunoSEQ software (Adaptive Biotechnologies) and ImmunoMap. *P* < 0.05 was considered statistically significant. Data are presented as mean ± SEM except for Figure 4B (median ± SEM).

---

**Supplementary Figure S1.**
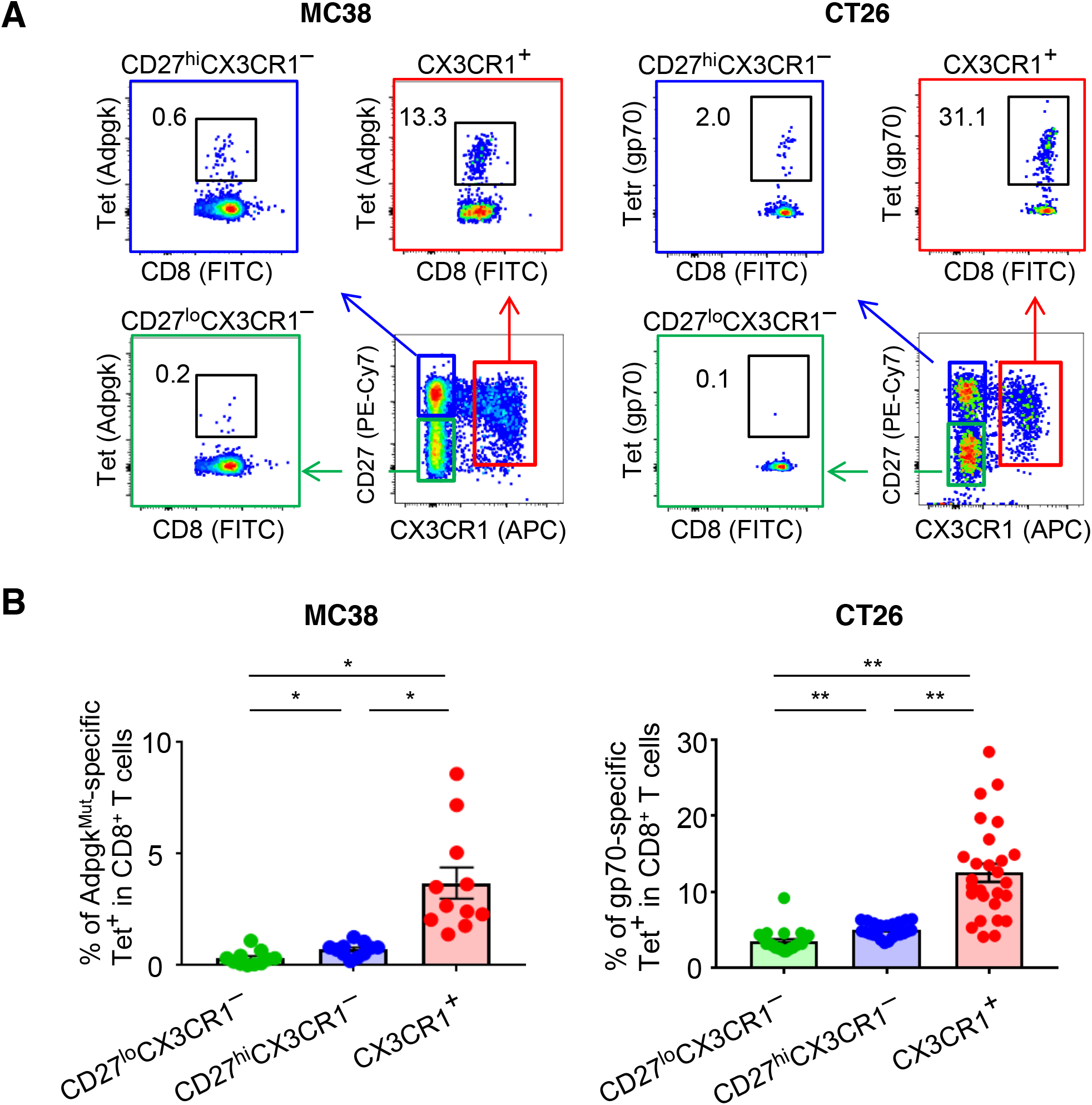
Related to Figure 2. **(A)** Representative flow cytometric plots showing the frequency of tetramer (Tet)^+^ CD8^+^ T cells in the CD27^lo^ CX3CR1^-^ (green), CD27^hi^ CX3CR1^-^ (blue), and CX3CR1^+^ (red) subsets in peripheral blood (PB) of MC38 (left) or CT26 (right) tumor-bearing mice treated with immune checkpoint inhibitors (ICI; anti-PD-L1 Ab and anti-CTLA-4 Ab) for 14 days; numbers denote percent Tet^+^ cells. We used a tetramer to detect CD8^+^ T cells specific for mutated Adpgk protein (Adpgk^Mut^) in MC38 and shared tumor-associated antigen (TAA), gp70 in CT26 tumor models **(B)** Frequency of Tet^+^ T cells in 3 subsets of PB CD8^+^ T cells at baseline in MC38 (left) and CT26 (right) tumor-bearing mice. n=3 experiments pooled. * *P*<0.005, \*\**P*<0.0001 by Mann-Whitney *U*-test. Values are mean ± SEM.

**Supplementary Figure S2.**
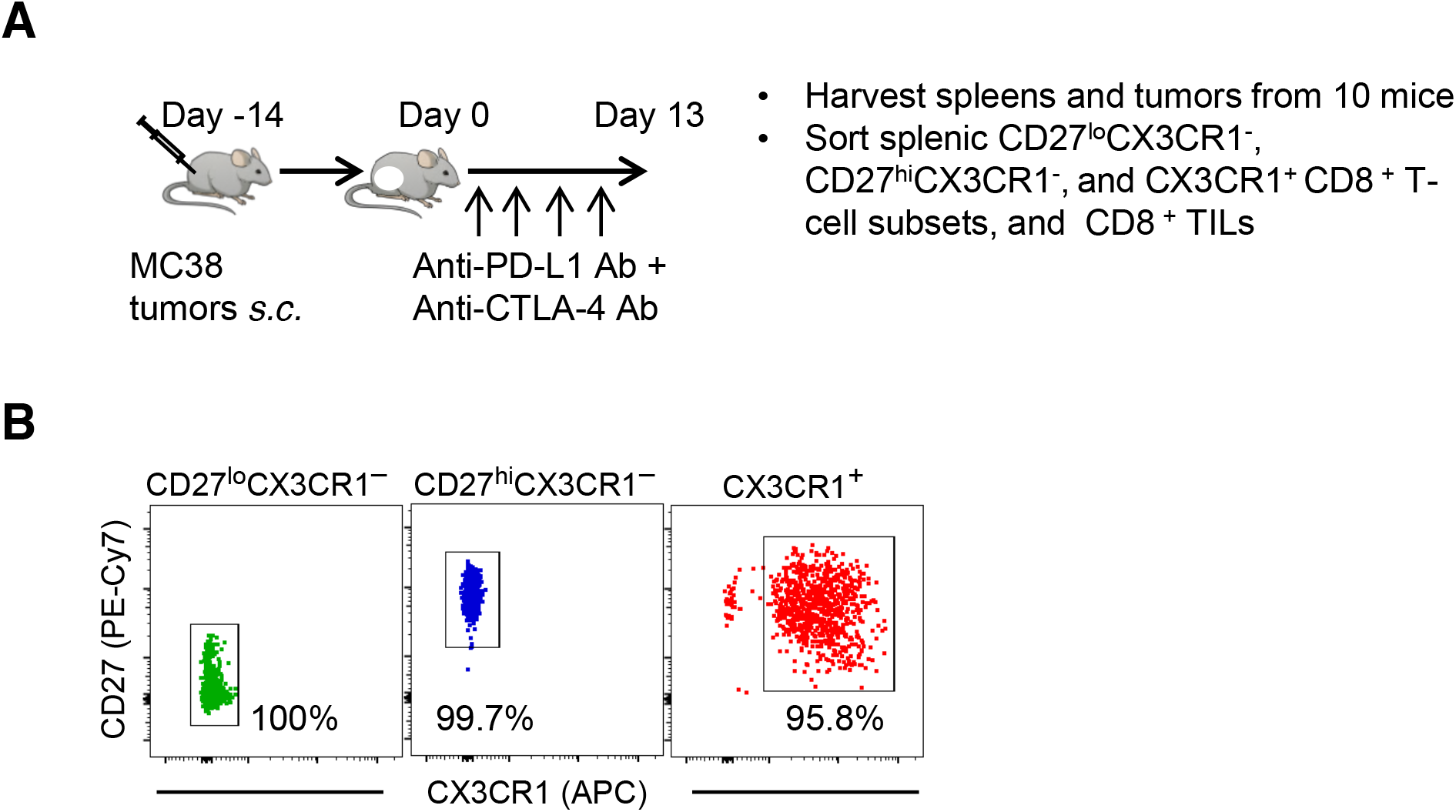
Related to Figure 3. (**A**) Experimental scheme of treatment with immune checkpoint inhibitors (ICI), and isolation of three subsets of splenic CD8^+^ T cells and CD8^+^ TILs for TCR repertoire and clonality analysis for figure 3, supplementary figures S3 - S5, and supplementary table S1. (**B**) Representative flow cytometric plots showing the frequency of splenic CD27^lo^ CX3CR1^-^ (green), CD27^hi^ CX3CR1^-^ (blue), and CX3CR1^+^ (red) CD8^+^ T cells after flow-sort.

**Supplementary Figure S3.**
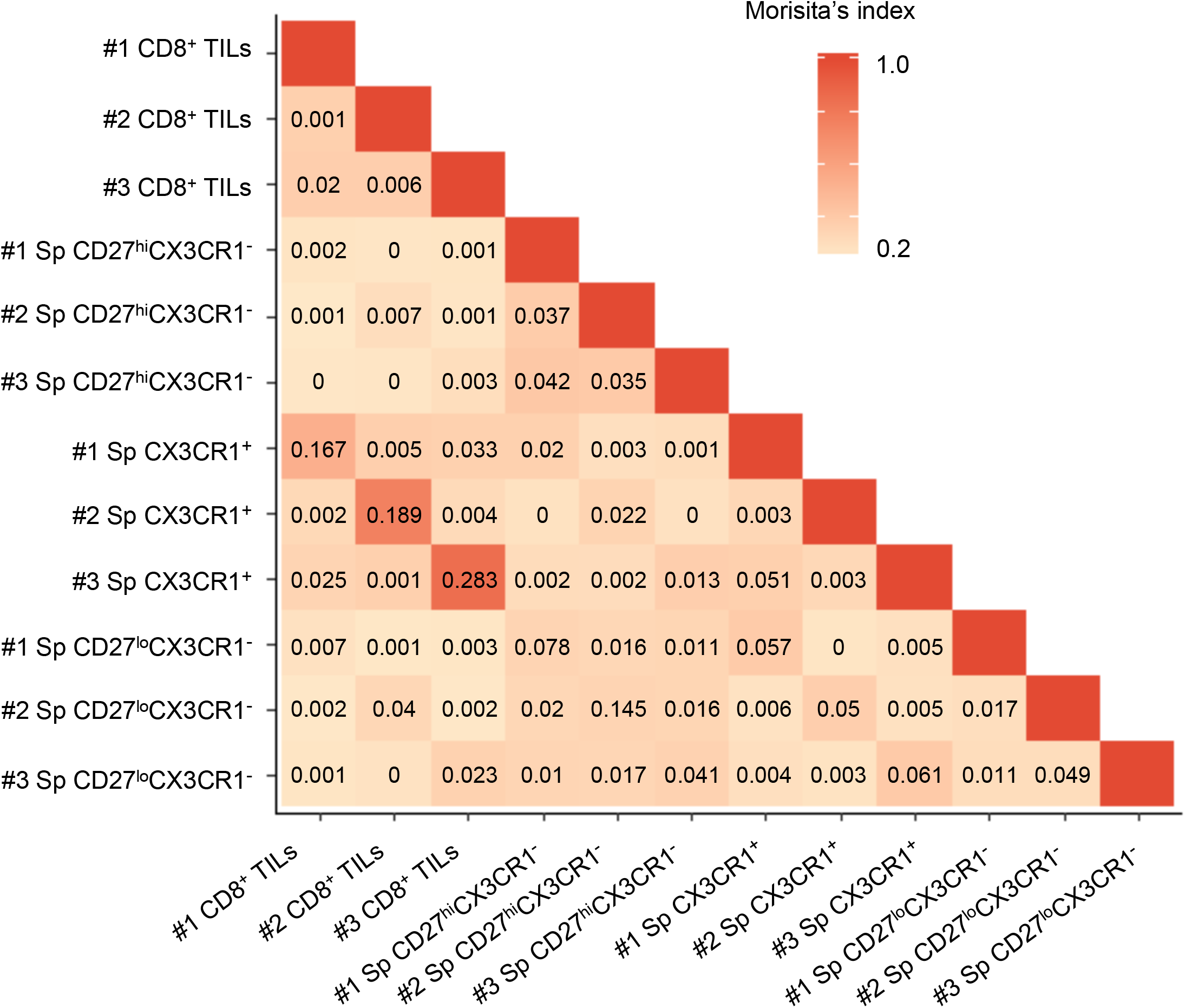
Related to Figure 3A. Pairwise TCRβ repertoire overlaps among different samples from three independent experiments (#1 - #3). Numbers denote TCR repertoire overlap by Morisita’s index.

**Supplementary Figure S4.**
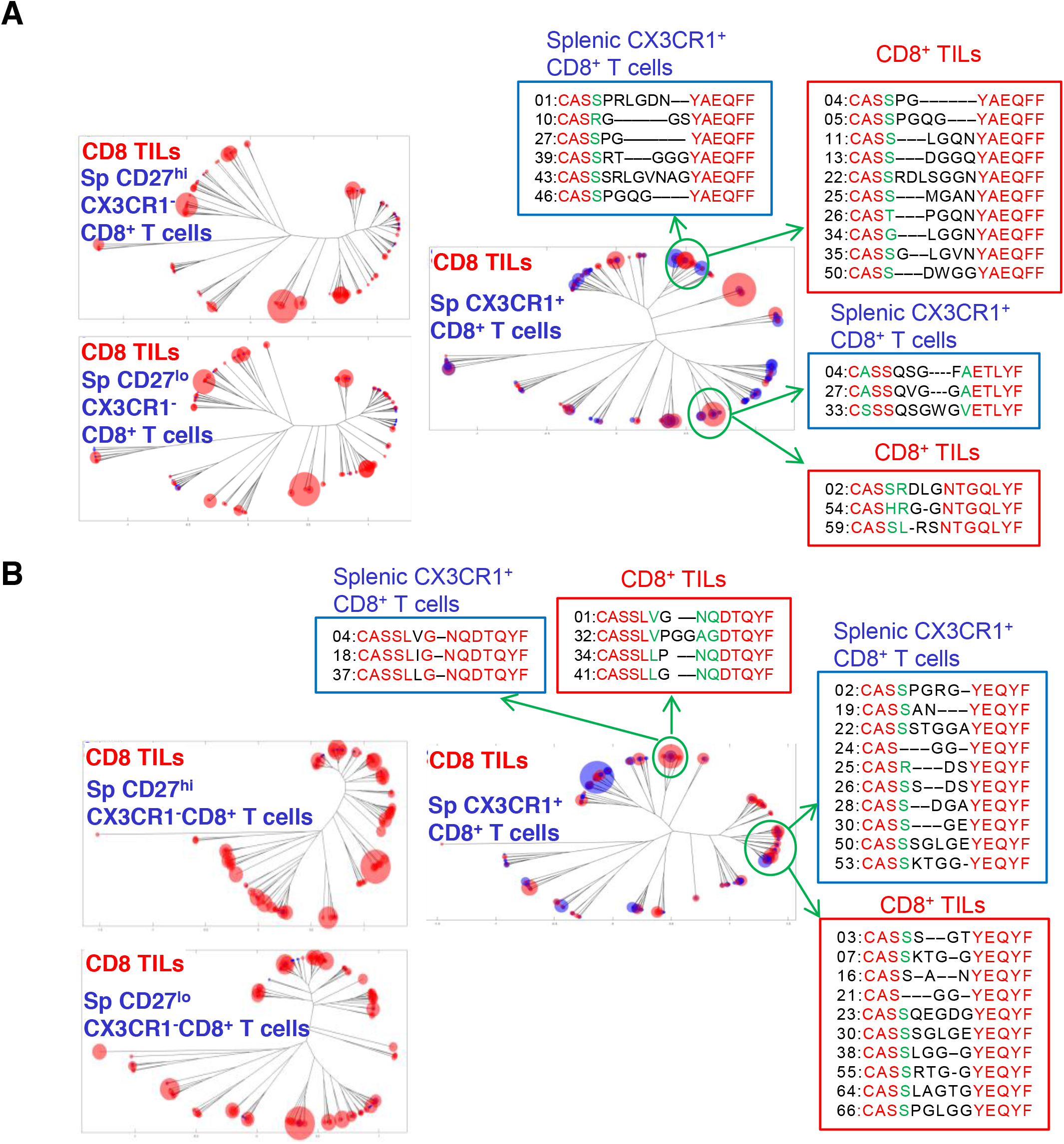
Related to Figure 3C and Supplementary Table S1. Representative overlapped weighted TCR repertoire dendrograms by ImmunoMap analysis between three subsets of splenic CD8^+^ T cells (blue) and CD8^+^ TILs (red) from two independent experiments (A and B). Dominant motif analysis clusters homologous sequences and selects for clusters contributing to significant proportion of the response. Two dominant motifs are shown representing highly represented structural motifs in each experiment. The numbers denote the ranking of the frequency within the subset. (red = fully conserved amino acids (AA), green = semi-conserved AA, black = non-conserved AA)

**Supplementary Table S1.** Demographic and clinical characteristics of lung cancer patients on this study

| Patient characteristics | *n* = 36 |
| --- | --- |
| Median age (range) | 68 (49-89) |
| Sex, *n* (%) |  |
| Male | 14 (39%) |
| Female | 22 (61%) |
| Race, *n* (%) |  |
| Caucasian | 34 (94%) |
| African-American | 2 (6%) |
| ECOG PS |  |
| 0/1 | 33 (92%) |
| History of smoking |  |
| Never | 4 (11%) |
| Former | 22 (61%) |
| Current | 10 (28%) |
| Histology, *n* (%) |  |
| Adenocarcinoma | 22 (61%) |
| Squamous cell carcinoma | 13 (36%) |
| Non-small cell lung cancer with giant features | 1 (3%) |
| Stage at diagnosis, *n* (%) |  |
| II-III | 9 (25%) |
| IV | 27 (75%) |
| Prior lung surgery, *n* (%) | 6 (17%) |
| Prior chemotherapy, *n* (%) |  |
| One line | 5 (14%) |
| Two lines | 2 (6%) |
| Prior targeted therapy, *n* (%) |  |
| Osimertinib and Erlotinib (EGFR) | 2 (6%) |
| Dabrafenib/Trametinib (BRAF V600E) | 1 (3%) |
| Prior radiation, *n* (%) |  |
| Thoracic radiation | 15 (42%) |
| Bone radiation | 3 (8%) |
| Gamma knife stereotactic radiosurgery | 7 (19%) |
| Known brain metastases, *n* (%) | 7 (19%) |
| Study drug, *n* (%) |  |
| Nivolumab | 2 (6%) |
| Pembrolizumab | 34 (94%) |
| Best disease response at 12 weeks, *n* (%) |  |
| Complete Response (CR) | 0 (0%) |
| Partial response (PR) | 13 (36%) |
| Stable disease (SD) | 14 (39%) |
| Progressive disease (PD) | 9 (25%) |

**Supplementary Table S2.** Marker Performance, Related to Figure 4B.

| PB CX3CR1<sup>+</sup>CD8<sup>+</sup> T cells | Cut-point | Specificity | Sensitivity | PPV | NPV |
| --- | --- | --- | --- | --- | --- |
| Max % change by 3 weeks<br>(n=27) | 1.74 | 0.65 | 0.80 | 0.57 | 0.85 |
| Max % change by 6 weeks<br>(n=36) | 21.19 | 0.87 | 0.62 | 0.73 | 0.80 |
| Max % change by 9 weeks<br>(n=36) | 15.49 | 0.74 | 1.00 | 0.68 | 1.00 |
| Max % change by 12 weeks<br>(n=36) | 19.62 | 0.71 | 0.92 | 0.80 | 0.95 |

Maximal percent change of the CX3CR1<sup>+</sup> subset in PB CD8<sup>+</sup> T cells at 3, 6, 9 and 12-weeks relative to baseline was evaluated. Cut-points for discriminating between responders and non-responders were obtained using the Youden's index criterion.
Abbreviations: PB, peripheral blood; Max, maximal; PPV, positive predictive value; NPV, negative predictive value.

**Supplementary Table S3.** Prediction performance for study biomarkers, Related to Figure 4E.

| | Baseline CD8<sup>+</sup> tumor-infiltrating lymphocytes (low/high vs. no/minimal infiltration) (n=24) | Baseline tumor mutational burden ($\geq 10.0/\text{Mb}$ vs. $<10.0/\text{Mb}$) (n=22) |
| --- | --- | --- |
| PPV | 33.3% (5/15) | 20.0% (1/5) |
| NPV | 55.6% (5/9) | 52.9% (9/17) |
| Sensitivity | 55.6% (5/9) | 11.1% (1/9) |
| Specificity | 33.3% (5/15) | 69.2% (9/13) |
| Accuracy | 41.7% (10/24) | 45.5% (10/22) |

Abbreviations: PPV, positive predictive value; NPV, negative predictive value

